# Red blood cell-tumour cell interactions promote tumour cell progression

**DOI:** 10.1101/2024.08.02.606121

**Authors:** Thais Pereira-Veiga, Celso Yáñez-Gómez, Aleksi Pekkarinen, Carmen Abuín, Christine Blechner, Miriam González-Conde, Alexander T. Bauer, Sabine Vidal-y-Sy, Ayham Moustafa, Bente Siebels, Ana B. Dávila-Ibáñez, Pablo Cabezas-Sainz, Maider Santos, Laura Sánchez, Joao Gorgulho, Julian Götze, Kira Meißner, Juan Cueva, Patricia Palacios, Alexia Cortegoso, Teresa Curiel, Carmela Rodríguez, Marta Carmona, Sven Peine, Milena Schmidt, Nadine Heuer-Olewinski, Martin Reck, Mustafa Abdo, Rafael López-López, Sabine Windhorst, Klaus Pantel, Harriet Wikman, Clotilde Costa

**Affiliations:** Department of Tumor Biology, University Medical Center Hamburg-Eppendorf, Hamburg, Germany; Translational Medical Oncology Group, Oncomet, Health Research Institute of Santiago de Compostela (IDIS), Santiago de Compostela, Spain; Department of Biochemistry and Signal Transduction, University Medical Center Hamburg-Eppendorf, Hamburg, Germany; Department of Dermatology and Venereology, University Medical Center Hamburg-Eppendorf, Hamburg, Germany; Section Mass Spectrometry and Proteomics, University Medical Center Hamburg-Eppendorf, Hamburg, Germany; CIBERONC, Centro de Investigación Biomédica en Red Cáncer, Madrid, Spain; Zoology, Genetics and Physical Anthropology Department, Universidade de Santiago de Compostela, Campus de Lugo, Lugo, Spain; Department of Oncology, Hematology and Bone Marrow Transplantation with Section of Pneumology, University Medical Center Hamburg-Eppendorf, Hamburg, Germany; University Clinical Hospital of Santiago (CHUS/SERGAS), Santiago de Compostela, Spain; Institute of Transfusion Medicine, University Medical Center Hamburg-Eppendorf, Hamburg, Germany; LungenClinic Grosshansdorf, Airway Research Center North, German Center for Lung Research, Grosshansdorf, Germany

**Keywords:** red blood cells, RBCs, breast cancer, lung cancer, circulating tumour cells, liquid biopsy, actin remodelling

## Abstract

One critical step in the metastatic cascade is the survival of circulating tumour cells (CTCs) within the bloodstream. While numerous interactions between CTCs and various hematopoietic cells have been described, the role of red blood cells (RBCs) in this process remains underexplored. This study investigates the interactions between tumour cells and RBCs from breast and lung cancer patients, revealing significant phenotypic and functional changes in the tumour cells, unlike when the contact is with RBCs from healthy donors*. In vitro* co-culture of cancer cell lines with RBCs from metastatic cancer patients resulted in increased tumour cell attachment accompanied by morphological changes. Additionally, RBCs-primed tumour cells showed increased adhesion and disruption of the endothelial barrier *in vitro* and increased invasiveness both *in vitro* and *in vivo*. Transcriptomic analysis showed that RBCs from metastatic breast cancer patients induce significant gene expression changes, notably upregulating *PAK4*, which enhances migration and epithelial-mesenchymal transition. PAK4 inhibition reduced these effects. Proteomic studies revealed substantial remodelling, including actin-related changes and the accumulation of VASP at cell edges, promoting directional migration. Clinically, higher RBC distribution width (RDW) in metastatic breast cancer patients is associated with increased CTC counts and worse outcome. This study highlights the previously unrecognized role of RBCs in promoting metastatic behaviours in cancer cells and suggests potential therapeutic targets, such as PAK4, to counteract these effects. Further exploration of RBCs-tumour cell interactions could provide new insights into metastatic mechanisms and improve cancer prognosis and treatment strategies.

**Key Points:**

- This study reveals the previously unknown role of RBCs in enhancing tumour cell invasiveness and metastatic potential.
- Tumour cells undergo significant phenotypic and functional changes after contact with RBCs from cancer patients.

## Introduction

Metastasis stands as the leading cause of cancer-related deaths^1^. Although extensive efforts have been made to investigate the metastatic cascade, significant gaps remain in our comprehensive understanding of the entire process. The metastatic cascade is an intricate sequence of events that cancer cells go through to move from the primary tumour to other locations within the body. It begins with local invasion, entering nearby tissues and blood vessels. Subsequently, these cells intravasate into the bloodstream or lymphatic system, becoming circulating tumour cells (CTCs), and travel to distant organs. In the blood, CTCs encounter significant challenges related to their survival in circulation and ability to extravasate to new metastatic sites^2^. Numerous important interactions between CTCs and blood components such as platelets, neutrophils, monocytes, or endothelial cells are crucial in the process. Many of these interactions are permissive or even necessary for CTCs survival in the bloodstream^2^. Surprisingly, the role of the most abundant component of the blood, the red blood cells (RBCs), remains almost unaddressed so far. RBCs initially thought to only carry oxygen, are now recognized for their role in maintaining metabolic homeostasis and influencing cellular processes^3^. Furthermore, RBCs have been implicated in the pathophysiology of diseases such as diabetes, Alzheimer’s, multiple sclerosis, and rheumatoid arthritis^4–7^.

While the interaction between tumour cells and RBCs remains understudied, the literature suggests that RBCs can interact with and modulate other immune cells such as eosinophils and lymphocytes^8–10^. RBCs have membrane-bound proteoglycans and glycoproteins that bind cytokines, thereby modulating inflammatory processes and potentially triggering cytokine storms e.g. after blood transfusions^11,12^. Cytokine profile alterations and immune modulations in RBCs have also been observed upon exposure to cancer cells^12^. Moreover, studies using mouse models have demonstrated that RBCs and haemoglobin can act as endogenous danger signals, promoting the proliferation of breast and melanoma tumour cells, as well as recruiting and polarizing macrophages^13^. Additionally, Helwa *et al*. proposed that tumour cells may interact with RBCs, potentially through galectin-4, which although primarily expressed in gastrointestinal epithelial cells, has implications in various cancers^8^. Furthermore, we have recently observed that the presence of RBCs in CTCs short-term cultures is associated with poorer patient outcomes^14^ and we have also demonstrated that the protein profiles of RBCs in cancer patients are altered compared to cancer-free donors^15,16^.

Clinically, research has been focused on the association of between red blood cell distribution width (RDW) and cancer. RDW, a routinely measured parameter in the complete blood count, is often altered in cancer patients and has been proposed as a novel biomarker for the diagnosis and prognosis of different tumour types^17–19^. However, the underlying biology of this phenomenon remains poorly understood.

In the bloodstream, CTCs have the opportunity to interact directly with numerous RBCs. To explore the possible consequences of these interactions, we conducted both *in vitro* and *in vivo* experiments using breast and lung cancer cell lines as model systems. RBCs from breast and lung cancer patients were used to prime the tumour cells for these experiments. This study is the first to comprehensively investigate the potential connections and biological consequences of tumour cells and RBCs interactions.

## Methods

Full description of methods can be found in the supplemental Information.

### Clinical samples

RBCs were isolated from venous blood from a cohort of both non-metastatic (M0) and metastatic (M1) breast cancer (n= 62) and non-small cell lung cancer (NSCLC) (n= 58) patients (at baseline, before treatment) and a cohort of healthy donors (HD) (n= 54) (supplemental Table 1-2). The samples were collected at the University Hospital Complex of Santiago de Compostela (CHUS, Spain) and at the University Medical Center Hamburg-Eppendorf (UKE, Germany). Whole blood samples from HD were provided by the Department of Transfusion Medicine (UKE) or volunteers (CHUS). All patients and HD gave written informed consent. The study was conducted according to the guidelines of the Declaration of Helsinki and approved by the Ethics Committee of Galicia (2015/772) and by the Ärztekammer of Hamburg (PV-5392).

### Enrichment of CTCs from blood samples

The CellSearch^®^ system (Menarini) and the Parsortix^®^ microfluidic system (ANGLE) were used for CTC detection from 7.5mL EDTA tubes of peripheral blood from breast (n= 55) and NSCLC cancer patients (n= 68), respectively, following manufacturer recommendations^20,21^.

### RBC isolation and tumour cell priming

Blood samples were collected in EDTA tubes and RBCs were isolated according to previously established protocols within two hours of collection^15^. Human cancer cell lines were co-cultured (primed) for 24 h with RBCs (2.10^6^ RBCs / 1.10^5^ tumour cells) before the start of experiments. Non-primed cell lines were used as a control (Ctrl) in each experiment.

### Cell culture

MCF-7 (ATCC HTB-22), MDA-MB-231 (ATCC CRM-HTB-26) cell lines were grown in complete DMEM supplemented with 10% Foetal Calf Serum (FCS). H1975 (ATCC CRL-5908) and A549 (ATCC CRM-CCL-185) cell lines were grown in complete RPMI-1640 supplemented with 10% FCS. HUVEC (ATCC PCS-100-013) cells were grown in EGM™-2 Endothelial Cell Growth Medium-2 BulletKit™ (Lonza) and culture plates were pre-treated with 0.2% gelatine (Sigma).

### Live-cell imaging

For time-lapse recordings, RBCs and H1975 cells (1.10^6^ RBCs/3,000 tumour cells) were placed in ultra-low attachment plates and imaged using a Zeiss Apotome microscope (Carl Zeiss) with an incubator (37°C, 5% CO_2_). A 20x objective lens was used, and images were captured every 5 minutes for over 18 h (n=3/group).

### Electric cell-impedance sensing (ECIS)

Endothelial cell monolayer impedance was measured continuously at different frequencies (500-64,000Hz) using an ECIS 1600R instrument (Applied BioPhysics, Inc.) as previously described^22^. HUVECs were grown to confluence and after 24 h, half of the cell culture medium was replaced by medium containing 1.10^4^ H1975 cells (non-primed, HD, and M1 primed RBCs). Results are expressed as relative electrical resistance with a 95% confidence interval (95% CI) (n= 3/group, duplicates).

### *In vivo* Zebrafish experiments

Between 100-200 tumour cells were injected into the Duct of Cuvier of each fish embryo (n= 3/group, 33 fish injected/sample) following standard procedures^23^. One-day post-injection, a fluorescence stereomicroscope (Nikon AZ-100) was used to image tumour spread and proliferation in the caudal hematopoietic tissue. Quantifish software was used to perform the image analysis ^24^.

### RNA-Sequencing analysis and RT-qPCR

RNA was extracted using the AllPrep DNA/RNA Mini kit (Qiagen) from MDA-MB-231 cells non-primed, primed with M1 RBCs from breast cancer patients, HD RBCs and liposomes^25,26^, mimicking RBC membranes (n=3/group, triplicates). cDNA libraries were created from 1 µg DNase-treated RNA with the TruSeq Stranded Total RNA Globin kit (Illumina) and sequenced on the Illumina NovaSeq6000. RNAseq data has been deposited at GEO under accession GSE273783. Gene expression was analysed with TaqMan assays for selected genes (supplemental Table 3), normalized to control Δct (n=14/group, triplicates).

### PAK4 inhibition

MDA-MB-231 cell line was exposed for 4 h to 10-60 μM of the PAK4 inhibitor LCH-7749944 (Selleckchem) to study the minimum dose of PAK4 inhibitor that affects proliferation (n= 3/group, quadruplicates). Proliferation was assessed at 24, 48 and 72 h using MTT assay (ThermoFisher). A concentration of 20 μM was chosen for the *in vitro* assays based on IC_50_ calculations.

### Mass spectrometry

Mass spectrometry-based proteomic analyses were performed as previously described^27^ (n= 5/group, 4 replicates). LC-MS/MS data were searched with the Sequest algorithm integrated into the Proteome Discoverer software (V3.0.0.757, ThermoFisher) against a reviewed human database. Carbamidomethylation was set as a fixed modification for cysteine residues. A maximum number of two missing tryptic cleavages was set. A cut-off (FDR < 0.01) was set for peptide and protein identification. Quantification was performed using the Minora Algorithm, implemented in Proteome Discoverer. Data have been deposited to the ProteomeXchange Consortium via the PRIDE partner repository with the dataset identifier PXD053123.

### Statistical analysis

Statistical analysis was conducted using R Studio (v4.3.0) and GraphPad Prism (v8.0). Chi-Squared or Fisher tests assessed categorical variable associations. Mann-Whitney U and Kruskal-Wallis tests evaluated differences between two or more groups. Progression-free (PFS) and overall survival (OS) were analysed with Kaplan-Meier plots and log-rank tests. Significance was set at P < 0.05. RNAseq data used DESeq2 for Fold Change and nbinomWaldTest. Gene enrichment was analysed with ShinyGO^28^. Proteins in ≥ 3 samples were tested with Student’s T-test (FDR q < 0.05), and significant proteins were analysed using IPA (Qiagen) ^29^.

## Results

### RBCs from metastatic patients interact with tumour cells causing their phenotypic transformation

To evaluate the interaction of RBCs with tumour cells *in vitro*, we co-cultured breast cancer (MDA-MB-231, MCF-7) and lung cancer (H1975, A549) cell lines in suspension for 24 hours with RBCs isolated from patients with localized (M0) or metastatic (M1) breast cancer or NSCLC disease, as well as from HD. We observed that M1 RBCs interact frequently with tumour cells while HD RBCs do not (Figure 1A). Thus, we observed a significantly higher percentage of cancer patients whose RBCs interact with tumour cells compared to HD (supplemental Figure 1A). Next, we quantified the interaction of individual tumour cells with RBCs from cancer patients and HD. Highly significant differences were found in the interaction between tumour cells and M1 RBCs compared with HD RBCs (Figure 1B).

**Figure 1.**
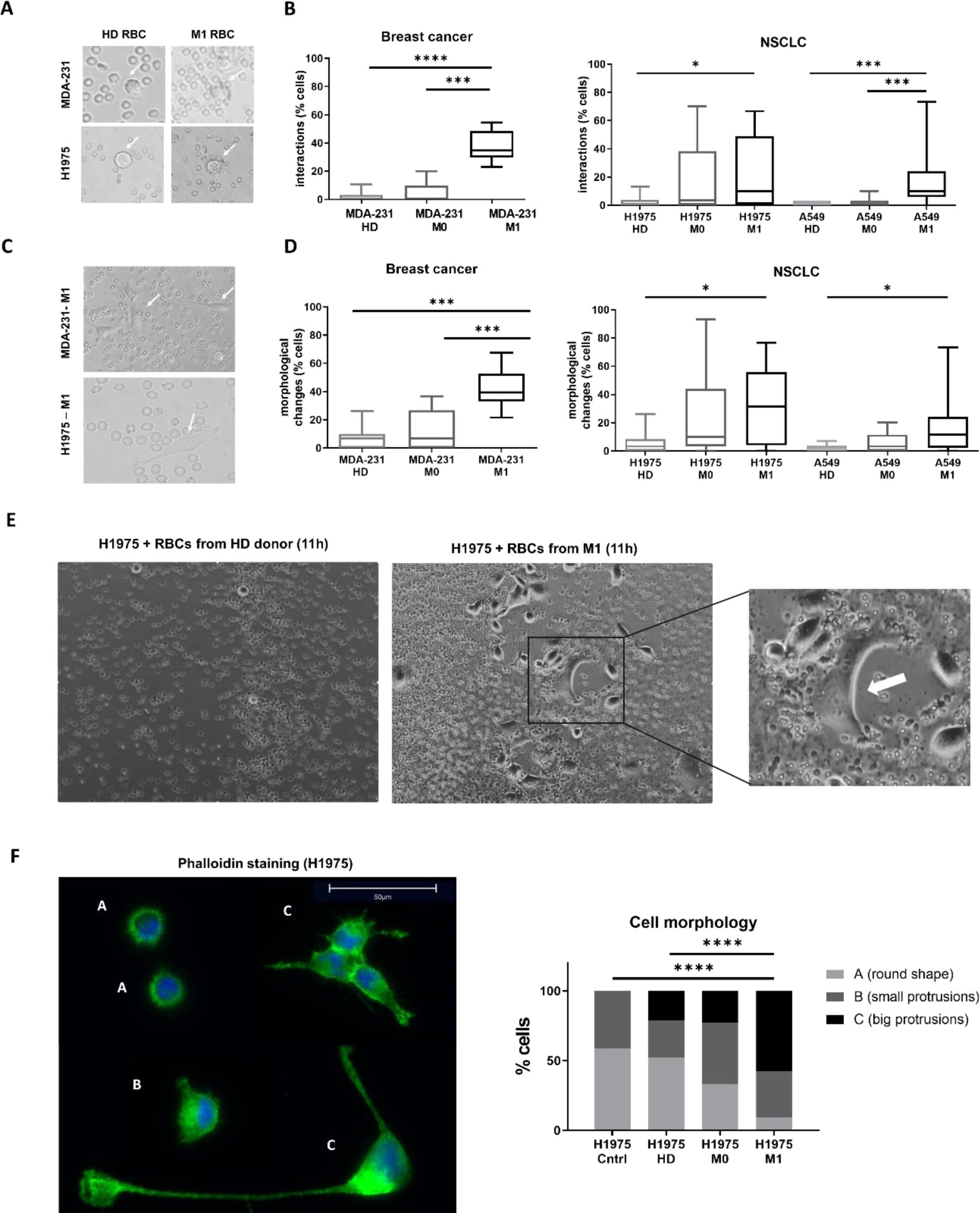
RBCs and tumour cells interact in suspension. **(A)** Exemplary pictures depicting the interaction between RBCs (from HD and M1) and tumour cells (MDA-MB-231 and H1975), showing a ring of RBCs from M1 patients attaching to a tumour cell, or no interaction for RBCs from HD (marked with a white arrow). **(B)** Boxplots showing the percentage of tumour cells (MDA-MB-231, H1975, and A549) interacting with RBCs from HD, M0 or M1 patients (n= 15 per group, 30 cells quantified/sample, triplicates). **(C)** Exemplary pictures depicting the change of morphology and adhesion of tumour cells under ultra-low attachment conditions. **(D)** Percentage of cells showing changes in morphology when MDA-MB-231 or H1975 are co-cultured with RBCs from M0 or M1 cancer patients compared to HD (n= 10-15 per group, 30 cells quantified/sample, triplicates). **(E)** Representative time-lapse images after 11 hours of co-culture with H1975 and RBCs from either a HD or a M1 NSCLC patient. The white arrow indicates the formation of lamellipodia-like structures (n= 3 per group). **(F)** Representative immunofluorescence images of morphological changes visualized by Alexa-fluor488-phalloidin staining of H1975 cells primed with RBCs (left panel) and quantification of type of morphological changes (right panel) (n= 5 per group, triplicates). * P < 0.05, ** P < 0.01, *** P < 0.001.

After co-culturing tumour cell lines with M1 RBCs in suspension, the tumour cells significantly more frequently attach to the bottom of the plates and exhibit morphological changes compared with those primed with HD RBCs (Figure 1C). This change was characterized by a transition from a rounded shape to an adherent growth form, exhibiting protrusions. To further investigate this observation, we conducted a 22-hour *in vitro* time-lapse live-cell imaging assay of H1975 cells with M1 or HD RBCs in ultra-low attachment conditions. Consistent with our previous data, H1975 cells exhibited major phenotypic changes after 12 hours when co-cultured with M1 RBCs, but not when co-cultured with HD RBCs. Representative images from the time-lapse live-cell imaging are shown in Figure 1E (complete video available in Video 1-2).

Based on the previous observation that cells co-cultured with M1 RBCs formed long protrusions, phalloidin staining was performed to visualize actin structures and identify different cell morphologies. For this aim, H1975 cells were primed under adherent conditions with HD, M0 or M1 RBCs. Tumour cells with large protrusions were significantly more frequently observed in the cells primed with M1 RBCs, while round shapes without protrusions were more common in cells primed with HD RBCs (Figure 1F). This indicates that priming tumour cell lines with RBCs from metastatic patients induces large actin rearrangement, favouring the formation of cells with numerous large protrusions.

### Patient-derived RBCs promote the tumour cell metastatic potential

Considering the significant impact of M1 RBCs on tumour cell phenotype, our next goal was to investigate whether priming human cancer cell lines with RBCs affects their functional abilities, including migration, adhesion, proliferation, and invasion—key steps in metastasis.

First, we studied whether RBCs from metastatic patients could affect tumour cell proliferation and migration. MDA-MB-231 and H1975 cells were primed with M1, M0 and HD RBCs. While no significant differences were found in the proliferation rates among the different conditions (supplemental Figure 1B), both cell lines showed significantly increased migration capability after 24 hours of priming with M1 RBCs compared to HD (Figure 2A). Next, we evaluated whether direct contact between the tumour cells and RBCs is necessary to enhance the migratory capacity of the tumour cells. For this, RBCs were seeded on the lower part of the transwell plate, hindering direct contact with tumour cells. In MDA-MB-231 cells, RBCs from metastatic breast cancer patients affected migration similarly to a direct contact setting priming, showing higher migration compared to non-primed cells. However, in H1975, although there is a trend towards a higher migration potential on metastatic RBCs, no significant difference was observed (Figure 2B). Notably, a migratory phenotype characterized by membrane protrusions resembling lamellipodia was detected mainly in the presence of M1 RBCs (supplemental Figure 1C).

**Figure 2.**
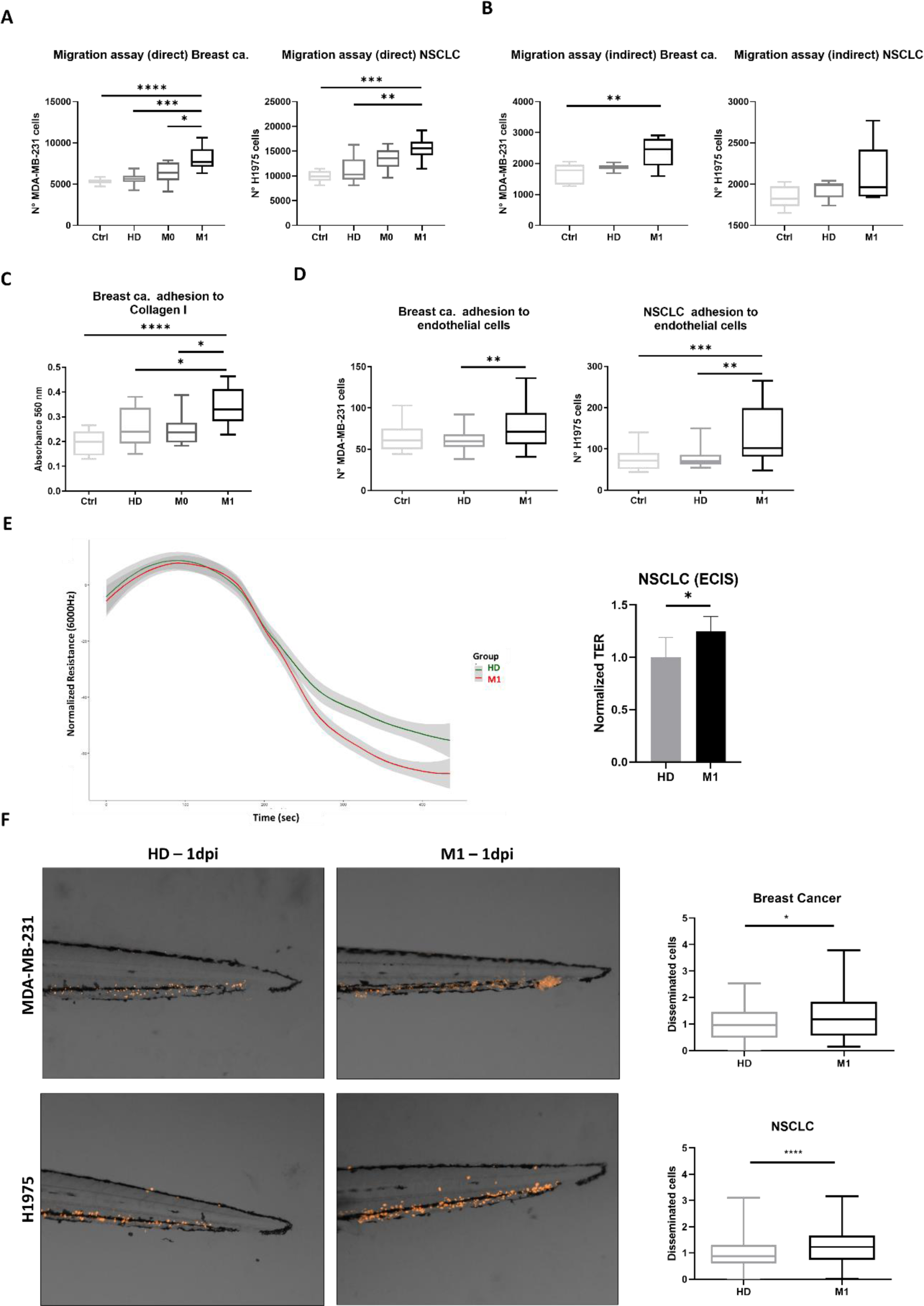
Functional evaluation of tumour cells after RBCs contact. **(A)** Boxplots depicting tumour cell transwell migration after priming (direct interaction) with RBC from HD, M0 or M1 for breast cancer (MDA-MB-231, left panel) and NSCLC cells (H1975, right panel) (n= 5 per group, triplicates). **(B)** Boxplot depicting tumour cell transwell migration with RBCs from HD and M1 in the bottom well (indirect interaction) for MDA-MB-231 (left panel) and H1975 (right panel) (n= 5 per group, triplicates). **(C)** Absorbance values reflecting the adhesion to collagen I by MDA-MB-231 after priming with RBCs from HD, M0 or M1 (n= 5 per group, triplicates). A negative control condition without priming with RBCs was included (Ctrl, n= 5). **(D)** Graph representing the MDA-MB-231 and H1975 cell counts (primed with RBS from HD and M1 or non-primed tumour cells (n=5 per group, triplicates) adhered to endothelial cells (HUVEC). **(E)** Results from the Electric Cell-substrate Impedance Sensing (ECIS) assay showing an increased disruption of the endothelial barrier function in the presence of H1975 cells primed with RBCs from M1 NSCLC patients compared to HD (n=3 per group, duplicates). Boxplot of the normalized transendothelial resistance (TER) of H1975 cells primed either with HD or M1 RBCs. **(F)** Representative images of tumour cell dissemination in the tail region of the zebrafish embryos injected with either H1975 or MDA-MB-231 cancer cell lines primed with RBCs from metastatic patients or HD, at 1dpi (n_total_ = 75 embryos per group, triplicates; survival MDA-MB-231 HD = 94, M1 = 75; survival H1975 HD = 162, M1 = 176). The main images are a superposition of a fluorescence image and a bright field image of the same embryo. Quantification of disseminated tumour cells on the zebrafish embryos tails at 1dpi. * P < 0.05, ** P < 0.01, *** P < 0.001.

Based on the observed tumour cell adhesion on ultra-low attachment surfaces, we assessed the adhesion of MDA-MB-231 cells to a collagen I matrix after priming with RBCs. As shown in Figure 2C, priming with M1 RBCs significantly increased adhesion compared to HD. Similarly, we found that the priming of both MDA-MB-231 and H1975 cells with M1 RBCs significantly enhanced their adhesion also to endothelial cells (HUVEC) *in vitro* (Figure 2D). Further analyses using the Electric Cell-substrate Impedance Sensing (ECIS) assay was used to determine the endothelial barrier function in the presence of tumour cells primed with M1 or M0 RBCs. This experiment revealed a 25% increase in the disruption of the endothelial barrier function induced by H1975 cells primed with M1 RBCs compared to HD RBCs (Figure 2E). A parallel assay with only RBCs from the different conditions was carried out, to discard background effect (data not shown).

To further validate our findings *in vivo*, zebrafish experiments were performed. MDA-MB-231 and H1975 cells primed with M1 or HD RBCs were injected into the Duct of Cuvier of zebrafish embryos. Tumour cells primed with M1 RBCs showed a significant dissemination enhancement to the caudal region (Figure 2F).

### Metastatic RBCs priming induces large gene expression changes and enhances breast tumour cell migration and adhesion via *PAK4*

Given previous findings of the impact of patient-derived RBCs on the phenotype and behaviour of tumour cells, we sought to investigate their effect on the gene expression profiles of tumour cells. Whole transcriptome sequencing revealed significant alterations in gene expression profiles between MDA-MB-231 cells primed with HD or M1 RBCs (Figure 3A-B) and with controls (supplemental Figure 1D). In total, 91 genes were significantly upregulated while 75 genes were downregulated between the cell lines primed with either M1 or HD RBCs. Gene Ontology analysis highlighted substantial changes in adhesion-related biological processes and genes involved in actin filament contraction and assembly (Figure 3C).

**Figure 3.**
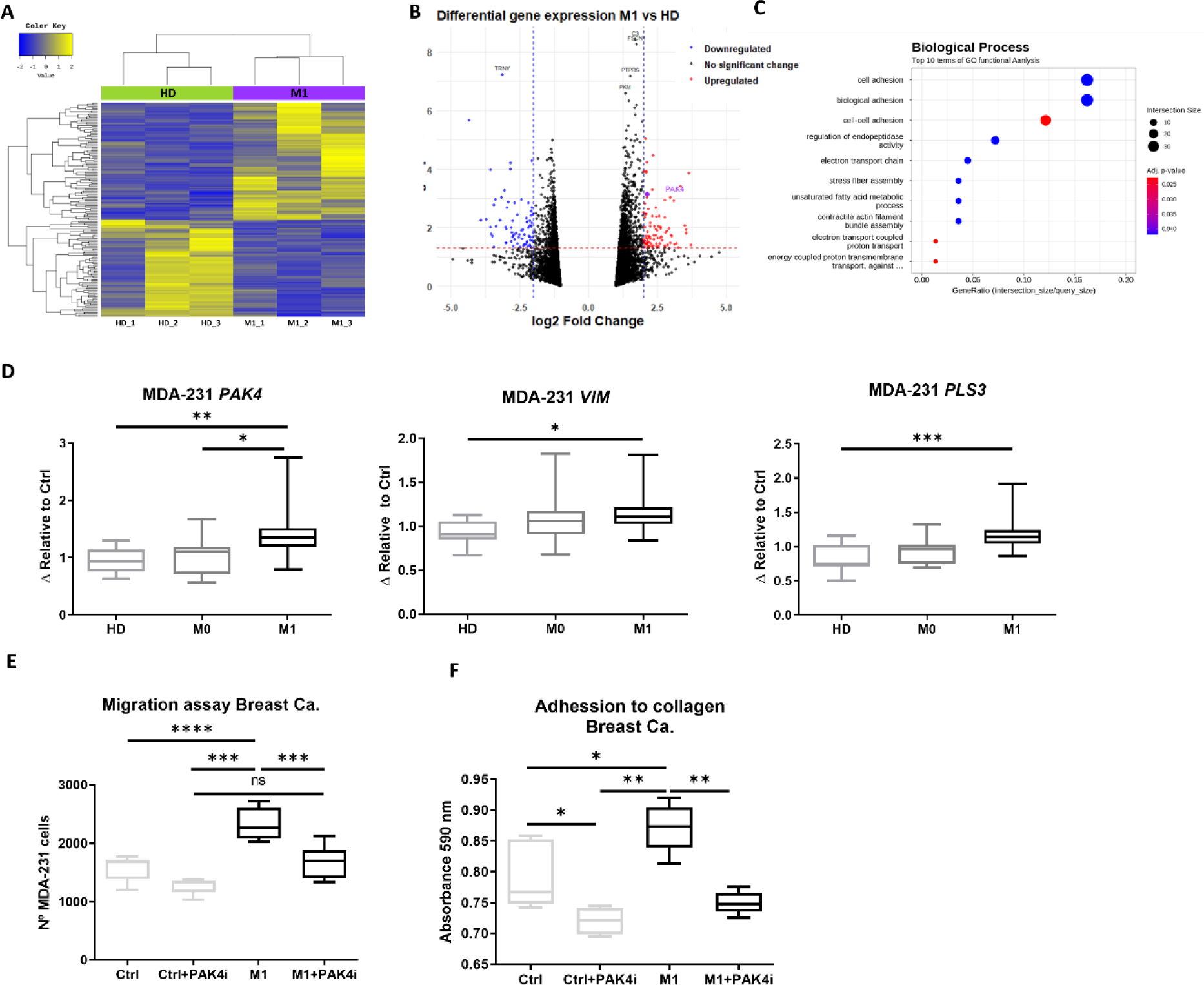
Transcriptomic and functional analysis of MDA-MB-231 cells primed with RBCs from M1 patients, HD and non-primed cells. **(A)** Heatmap showing results of the hierarchical clustering analysis of the significantly differentially expressed genes in MDA-MB-231 primed with RBCs from either HD (green) or M1 patients (purple) (n= 3 per group). **(B)** Volcano plot for the differentially expressed genes between HD and M1 samples. **(C)** Gene ontology analysis of the differentially expressed genes displaying the biological processes altered on MDA-MB-231 after co-cultivation with RBCs. (D) *PAK4, VIM* and *PLS3* gene expression was analysed by RT-qPCR, from MDA-MB-231 samples co-cultured with HD, M0 or M1 RBCs. Samples were relativized to *B2M* and normalized to Δct from negative control (non-primed cells) (n= 14 per group, triplicates). **(E)** Graph representing migration of MDA-MB-231 co-cultured with metastatic breast cancer RBCs or without RBCs in the presence or absence of PAK4 inhibitor (PAKi, LCH-7749944) (n= 5 per group, duplicates). **(F)** Graph representing adhesion to collagen I of MDA-MB-231 primed with metastatic breast cancer RBCs or without RBCs in the presence or absence of PAK4i (n= 5 per group, triplicates). * P < 0.05, ** P < 0.01, *** P < 0.001.

Among the genes with higher expression in the metastatic group (supplemental Table 4), *PAK4* was chosen for its role in survival, migration, and epithelial-mesenchymal transition (EMT). The expression changes were validated by RT-qPCR in tumour cells primed with a larger cohort of RBCs from cancer patients and HD. *PAK4* was overexpressed in MDA-MB-231 cells primed with M1 RBCs compared to cells primed with HD or M0 RBCs (Figure 3D). Further investigation into EMT-related genes in MDA-MB-231 cells revealed that cells co-cultured with M1 RBCs exhibited significantly higher levels of Vimentin (*VIM*) and Plastin 3 (*PLS3*) compared to HD RBCs (Figure 3D). To further investigate the role of PAK4 in the observed functional changes in MDA-MB-231 cells, we used a PAK4-specific inhibitor (LCH-7749944, PAK4i) (supplemental Figure 1E). PAK4 inhibition significantly reduced the migration capacity of MDA-MB-231 cells primed with M1 RBCs and non-primed cells. Importantly, no significant differences were found between non-primed cells and M1 RBCs-primed cells treated with PAK4i, indicating that the inhibitor reversed the enhanced migratory capacity of the cells (Figure 3E). Consistent with these findings, PAK4 inhibition in MDA-MB-231 cells primed with M1 RBCs reduced adhesion to collagen I levels to those comparable with non-primed cells (Figure 3F).

### RBCs from metastatic cancer patients induce global proteomic changes including actin remodelling in lung cancer cells

We investigated the global proteomic changes in H1975 cells primed with RBCs from NSCLC patients to identify the underlying processes and molecular targets influenced by the presence of RBCs. In total, 4414 proteins were identified, of which 626 proteins were significantly differentially abundant between HD, M0 and M1 RBCs primed cells (Figure 4A-B, supplemental Figure 2A). This indicates that there is a large remodelling of the proteome taking place in the tumour cells following contact with RBCs. We detected a higher differential abundance of 269 and lower differential abundance of 176 proteins in H1975 primed with M1 RBCs compared to HD RBCs. Among the top 20 differentially abundant proteins, we identified two EMT related proteins, KRT5 and CDH1 (supplemental Table 5), which were also validated by immunofluorescence staining (Figure 4C).

**Figure 4.**
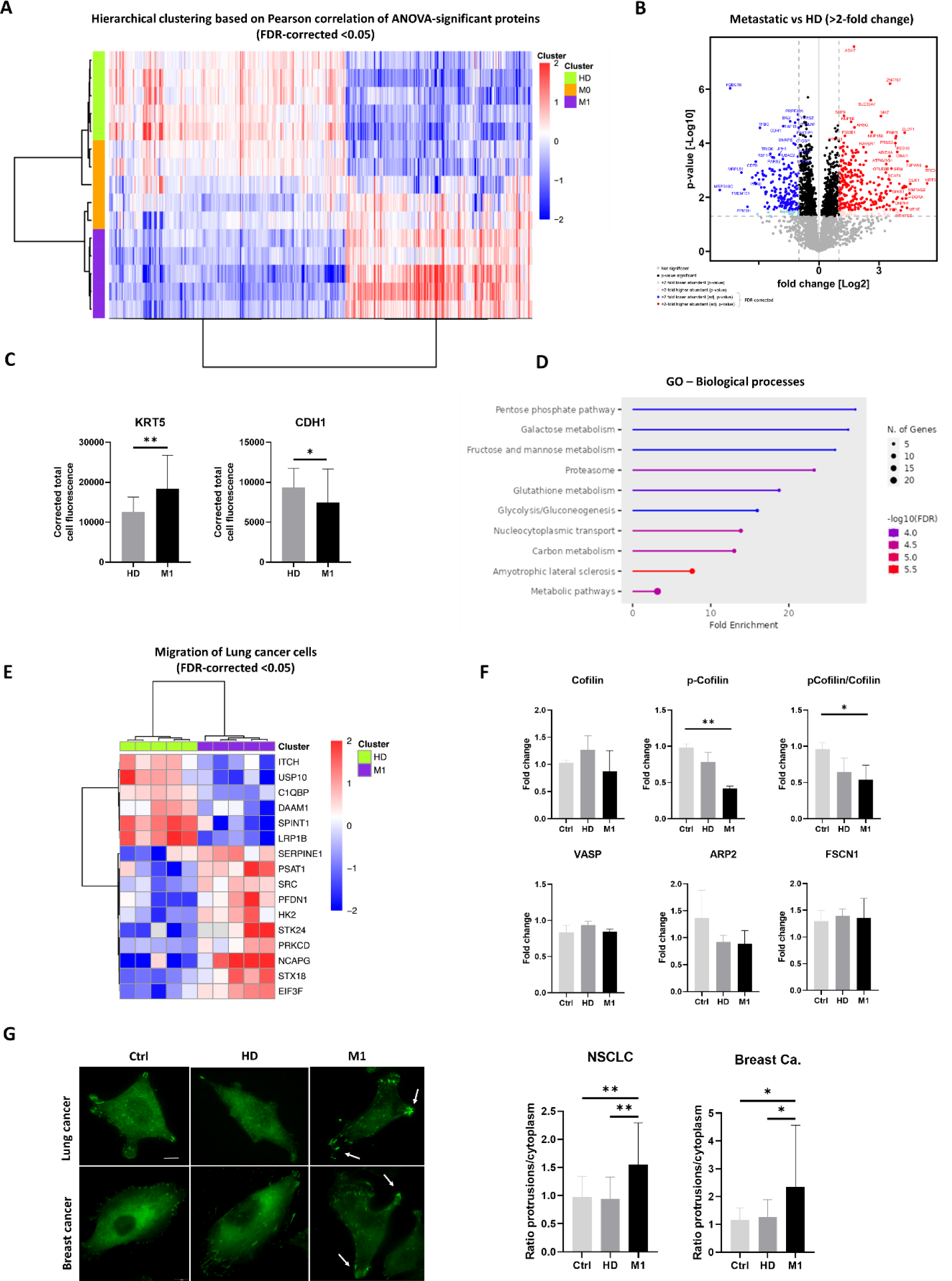
Proteomic analysis of tumour cells primed with RBCs from M1 and M0 patients, HD and non-primed cells. **(A)** Hierarchical clustering based on Pearson correlation of ANOVA-significant proteins of H1975 cell primed with RBCs from HD, M0 and M1 patients (n= 5 per group, quadruplicates). **(B)** Volcano plot for the differentially abundant proteins between M1 and HD samples. **(C)** Bar plot representing the quantification of the fluorescence intensity in H1975 cells stained with anti-KRT5 and anti-CDH1, post-priming with RBCs from HD or metastatic patients (n= 30 cells quantified/group, n= 3). **(D)** Gene ontology analysis of the differentially expressed proteins showing the biological processes affected on H1975 cells after priming with RBCs. **(E)** Hierarchical clustering of the differential abundance of the proteins from the dataset involved in migration of lung cancer cell lines, between H1975 cells primed either with HD or M1 RBCs (FDR < 0.05, Fold-change >2). **(F)** Bar plots representing the quantification of western blots for the proteins Cofilin, p-Cofilin, VASP, ARP2 and FSCN1 in H1975 primed with RBCs from HD or metastatic patients as well as non-primed cells (Ctrl) (n= 6 per group). **(G)** Representative images of H1975 cells and MDA-MB-231 cells, stained for the protein VASP, after priming with RBCs. The white arrows indicate the accumulation of VASP on the cellular protrusions (left panel). Quantification of the ratio of VASP-fluorescence intensity in protrusions/cytoplasm on H1975 and MDA-MB-231 cells stained with VASP after being primed with RBCs from HD or metastatic RBCS, as well as non-primed cells (n=30 cells quantified/group, n= 3) (right panel). * P < 0.05, ** P < 0.01, *** P < 0.001.

The gene ontology analysis revealed significant alterations in the enrichment of proteins related to pathways mainly involved in metabolic processes (Figure 4D). Further analyses using the IPA and z-score algorithm identified migration, cell movement, invasion and DNA repair to be significantly increased, while autophagy was predicted to be decreased. More specifically, migration of lung cancer cell lines was predicted to be increased with 16 proteins significantly differentially expressed (Figure 4E, supplemental Figure 2B-C). Furthermore, the activity of 11 upstream regulatory proteins was predicted to be activated, including up-stream Rab-like protein 6 (RABL6), Heat-shock factor 1 (HSF1), Epidermal grow factor (EGF) and ERBB2, which are involved in progression and development of several types of cancer (supplemental Figure 2D).

Since cytoskeletal dynamics play a crucial role in regulating cancer cell adhesion, migration, and invasion, we examined the activity and expression levels of different proteins involved in actin remodelling in more detail. We began by analysing with western blot the activity of Cofilin-1, an actin-severing protein and downstream effector of the Cdc42-PAK-LIMK pathway, whose activity is inhibited by LIMK-mediated phosphorylation at Ser3. The level of phosphorylated Cofilin (p-Cofilin) was significantly decreased in cancer cells treated with M1 RBCs, whereas the total Cofilin-1 protein level remained unchanged, leading to a significantly decreased ratio of p-Cofilin/Cofinin-1 (Figure 4F, supplemental Figure 2E). To determine if also proteins that promote F-actin elongation or bundling were altered in cells treated with RBCs from metastatic patients, we analysed the protein levels of Arp2, VASP and Fascin by Western blotting. Although we did not find significant differences in the total protein levels of these proteins between the control group and cells incubated with M1 RBCs (Figure 4F and supplemental Figure 2F), immunofluorescence analysis revealed a significant increase in the cellular localization of VASP at cellular protrusions in both lung and breast cancer cells (Figure 4G). This suggests that signals derived from RBCs of metastatic patients lead to the accumulation of VASP at the leading edge of cancer cells, promoting migratory capabilities towards a specific direction.

### Clinical implications of the interaction of RBCs from metastatic patients with CTCs

To analyse whether alterations in patients’ RBCs have clinical implications, we checked if CTCs count correlates with the RDW values and outcomes. For this, 7.5ml of blood from 55 metastatic breast cancer patients were assessed for CTCs enumeration by the CellSearch^®^ system at diagnosis of metastatic disease (before start of therapy). The mean number of CTCs was 101.8. Notably, 84.0% of patients had at least one CTC, with 51.4% of patients meeting the standard cut-off of ≥ 5 CTCs/7.5 ml blood^30^. Patients with high RDW (≥ 14.5) had significantly more CTCs, compared to patients with normal RDW (Figure 5A). Next, we performed a survival analysis based on RDW levels and found that high RDW was associated with worse PFS and OS (Figure 5B). Interestingly, the presence of ≥ 1 CTC was associated with worse OS but not with PFS (supplemental Figure 3A-D). The combination of high RDW and the presence of either ≥ 1 CTC or ≥ 5 CTC was associated with both worse PFS and OS, showing increased significance compared to RDW alone or CTCs alone (Figure 5C-D). Multivariate analysis indicated that both CellSearch^®^ and RDW values are independent parameters for PFS an OS prediction. The same analysis was performed on NSCLC patients (n=68, stage I-IV) with CTCs enumerated by Parsortix^®^. RDW alone was significantly associated with OS, however neither CTCs alone nor RDW in combination with CTCs was significant, probably due to the low number of CTCs on these samples (14.5% had ≥ 1 CTCs/7,5ml) (supplemental Figure 5E-G). We have previously reported that the presence of RBCs in short-time breast cancer CTC cultures is associated with worse outcomes^14^. Interestingly, we could also observe some direct contacts between RBCs and CTCs in these patient-derived cultures (Figure 5E-F).

**Figure 5.**
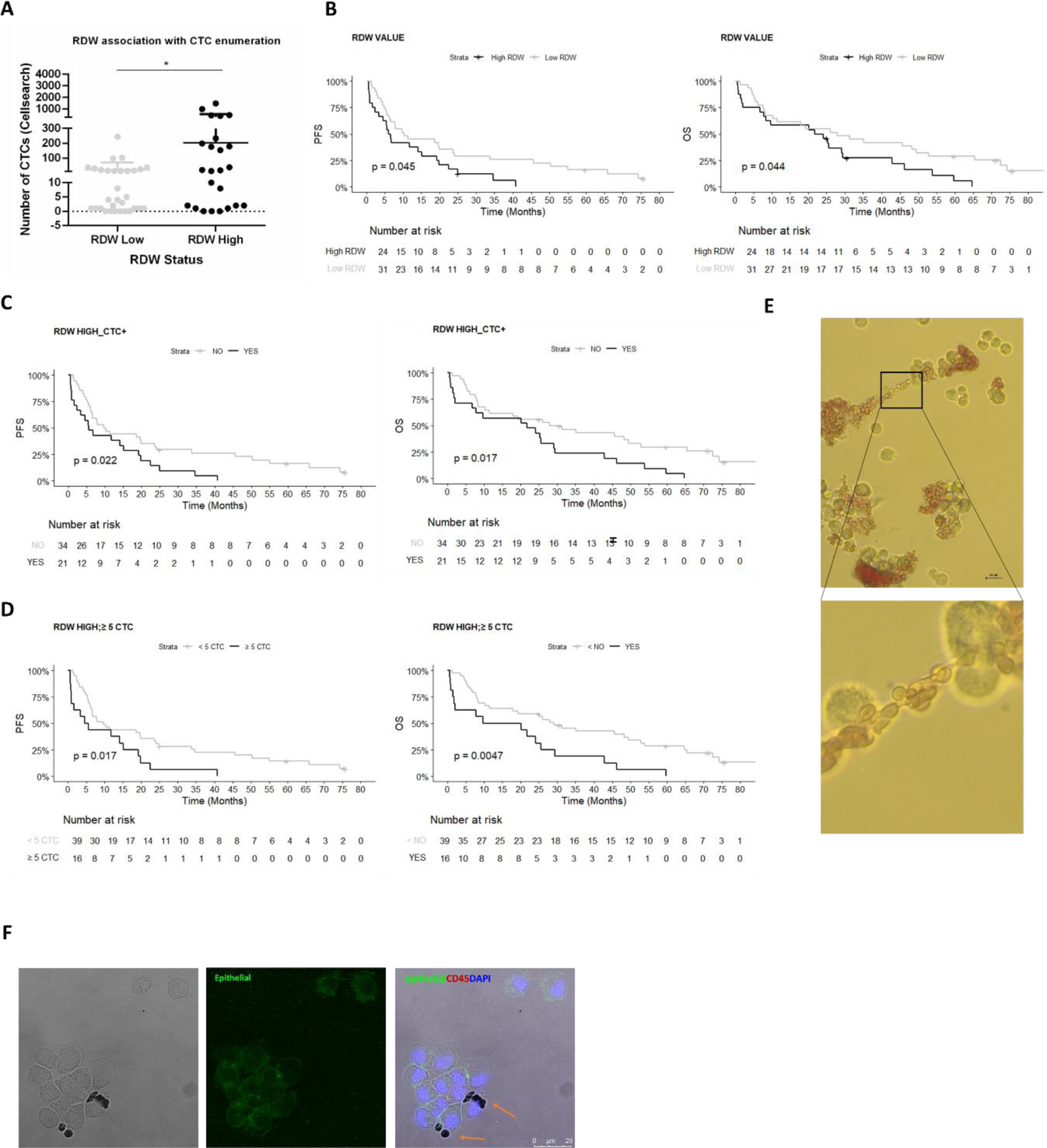
RBC interaction with CTCs. **(A)** Box plot of individual values representing the association of RDW status and CTCs number, determined by CellSearch^®^ from 7.5 mL of blood from metastatic breast cancer patients (n=55). **(B)** Kaplan–Meier plots of PFS and OS for high (black) or low (grey) RDW values (cut-off = 14.5) in breast cancer patients (median PFS: 6.38 vs 10.29 months, P = 0.04; median OS: 22.9 vs 27.8 months, P = 0.04) **(C)** Kaplan–Meier plots of PFS and OS for patients with a combination of high RDW and ≥ 1 CTC (RDW HIGH_CTC+, black) or not (grey) (RDW cut-off = 14.5) (median PFS: 5.56 vs 10.01 months, P = 0.022; median OS: 21.8 vs 27.8 months, P = 0.017) **(D)** Kaplan–Meier plots of PFS and OS for patients with a combination of high RDW and ≥ 5 CTC (RDW HIGH_≥ 5 CTC, black) or not (grey) (RDW cut-off = 14.5) (median PFS: 9.73 vs 5.08 months, P = 0.017; median OS: 29.1 vs 14.9 months, P = 0.0047) **(E)** Representative brightfield microscope images of CTCs isolated from a metastatic breast cancer patient in *ex vivo* culture with RBCs attached around them. **(F)** Representative images by confocal microscopy of immunofluorescence characterization of CTCs from a metastatic breast cancer patient after 10 days of culture. Immunofluorescence staining was performed using a combination of anti-human epithelial markers (EpCAM, CDH1, and PanCK, green), anti-CD45 (red) and DAPI (blue). RBCs are attached to the CTCs and are pointed with orange arrows. * P < 0.05.

## Discussion

Despite being a significant component of the blood, the role of RBCs has been largely overlooked in cancer and metastasis^2^. In this study, we explored the interactions between tumour cells and RBCs and their potential influence on the metastatic processes. Our results using tumour cell lines and RBCs from cancer patients and HD show that RBCs from metastatic breast and lung cancer patients closely interact with cancer cell lines *in vitro*, unlike RBCs from HD. This is, to our knowledge, the first study showing that this communication leads to profound metabolic, morphologic, and transcriptional changes in both lung and breast tumour cells, enhancing their metastatic potential and malignancy.

We could observe that cancer cells in the presence of RBCs, especially from metastatic cancer patients, change cell morphology, transitioning from a rounded shape to a more mesenchymal cell type with lamellipodia-like protrusions, key membrane structures involved in chemo-sensing and migration^31–33^ and commonly associated with increased metastatic potential^34^. Furthermore, we also observed that tumour cell lines show enhanced migratory and adhesive capabilities after having direct contact with RBCs from cancer patients. In addition, we also observed that tumour cells exposed to RBCs from metastatic patients were able to significantly disrupt the endothelial barrier, potentially facilitating trans-endothelial migration of CTCs in blood circulation. Furthermore, *in vivo* analyses using a zebrafish model confirm our findings showing an increased dissemination of tumour cells primed with RBCs from metastatic patients. These results suggest that in metastatic cancer patients, RBCs play a role in the modulation of tumour cells, rendering them more aggressive, possibly playing a pivotal role in the metastasis and progression of tumours.

Moreover, we observed large global effects on both the transcriptomic and proteomic levels, as demonstrated by RNAseq and MassSpec analysis, respectively. RNAseq analysis in breast cancer cells revealed enrichment of adhesion-related genes and pathways involved in actin filament assembly, consistent with our experimental findings. Among the genes affected, the upregulation of *PAK4, VIM* and *PLS3*—all pivotal in cytoskeletal organization, cell motility, and EMT^35^ was validated by RT-qPCR analysis in MDA-MB-231 cells primed with RBCs from metastatic breast cancer patients.

All three genes have been described to be upregulated in various cancer types, including breast cancer, and have been correlated with poorer prognosis in breast cancer patients^36–39^. Importantly, the use of a specific PAK4 inhibitor (LCH-7749944) effectively reversed the increased migration and collagen I adhesion observed in MDA-MB-231 cells following exposure to RBCs from metastatic breast cancer patients. This suggests that *PAK4* may indeed mediate the phenotypic and functional changes in breast cancer cells induced by these RBCs. Clinical validation of PAK4 inhibitors such as LCH-7749944 is promising, as they have shown efficacy in preclinical models and are currently being evaluated in clinical trials^40,41^. These inhibitors hold potential for therapeutic intervention aimed at mitigating the aggressive behaviour of breast cancer cells influenced by RBCs from metastatic patients.

Similar to findings in breast cancer, MassSpec analyses of H1975 cells primed with RBCs from HD or cancer patients revealed profound alterations in the proteomic composition of these tumour cells. Notably, while proteins like VIM and PLS3 showed no significant changes, other markers of EMT such as KRT5 and CDH1 were affected, suggesting shared pathways but distinct proteins as protagonists between breast cancer and NSCLC. IPA analyses revealed the enrichment of metabolic pathways and predicted increases in migration, cell movement, and invasion-related proteins in line with previous *in vitro* observations. Furthermore, altered cytoskeletal dynamics, highlighted by reduced phosphorylation and thereby activation of Cofilin-1 and increased VASP localization at cell protrusions, suggest mechanisms through which RBCs interactions may enhance actin turnover. These observations align with findings from Sidani *et al*., which demonstrated that modulation of Cofilin-1 expression induces phenotypic shifts from amoeboid to mesenchymal cancer cell types^42^. Reduced Cofilin-1 levels were furthermore associated with decreased free F-actin barbed ends and increased stability of directed Arp2/3-mediated actin branches. This mechanistic insight underscores how interactions with RBCs from metastatic patients may drive cytoskeletal rearrangements leading to a more mesenchymal phenotype with enhanced migration. Our findings suggest that RBCs, within an altered systemic environment, promote cancer progression by influencing the aggressiveness of tumour cells in numerous ways. Previously, we reported modifications in RBCs protein profiles in metastatic breast cancer patients^2^. Together with the results presented in this work, the data indicates a significant crosstalk between tumour cells and RBCs, leading to profound alterations in both entities, especially in metastatic conditions. Importantly, since mature RBCs lack nuclei and cannot synthesize new RNA, priming could potentially occur earlier in the bone marrow, possibly via paracrine interactions such as extracellular vesicles or secreted factors^43^.

Finally, patients with elevated RDW, a numerical measure of anisocytosis in RBCs^44^, exhibited worse outcomes as previously described^17,45,46^, particularly when CTCs were detected in the blood. Notably, RDW and CTCs represent independent variables providing distinct information about the patient’s condition; RDW reflects systemic health while CTCs indicate tumour burden^47–49^. Furthermore, their combined assessment could provide a more robust prognostic biomarker. However, these intriguing findings raise numerous questions for further investigations. Indeed, some limitations of our work, that should be covered in future studies, could include exploring the direct versus indirect roles of RBCs interactions involving larger sample sizes and identifying key regulatory mechanisms underlying RBCs-mediated effects in cancer progression.

In summary, this study uncovers a novel role for RBCs in enhancing tumour cell survival and invasiveness during metastasis. By revealing significant gene expression changes and protein remodelling, such as PAK4 upregulation and actin-related modifications, this research highlights mechanisms driving tumour cell migration and EMT. Clinically, we have shown that integrating RBCs parameters (RDW) and CTC numbers improves the prognosis prediction in metastatic breast cancer. These findings open new opportunities for understanding metastatic mechanisms.

## Supporting information

Supplemental data_Red blood cell-tumour cell interactions promote tumour cell progression

## Acknowledgements

The authors thank Dr. Antonio Virgilio Failla and Dr. Bernd Zobiak (UKE Microscopy Imaging Facility, University Medical Center Hamburg-Eppendorf) for their technical and scientific support. Also, the authors would like to thank the patients who participated in this study; all the personnel of the Oncology Service at the University Clinical Hospital of Santiago de Compostela for their help with patient care and sample management, especially Rosa Villaverde. This study was supported by grants from the Deutsche Forschungsgemeinschaft (DFG) (DFG-SPP 2084: µBONE, INST 337/15-1, INST 337/16-1, INST 152/837-1 and INST 152/947-1 FUGG) and Axencia Galega de Innovación (GAIN); Vicepresidencia Segunda e Consellería de Economía; Empresa e Innovación (IN853B 2018/03) and Foundation Spanish Association Against Cancer (IDEAS18108COST). C.Y.G. was supported by Axudas Predoutorais da Xunta de Galicia, and CC was supported by the Foundation Spanish Association Against Cancer (AECC, in Spanish) (INVES211437COST) and Instituto de Salud Carlos III (ISCiii) (CP23/0002).

## Authorship contribution

CC, TPV, CYG, HW and SW conceived the study. TPV, CYG, AP, CA, CB, SW, MGC, ATB, SV-y-S, PCS, MS, AM and BS performed and analysed experiments. KP, HW, TPV, CC, CYG, SW, ATB, PCS, RLL, BS and AM helped shaped the research and analysis. ADI, LS and SW contributed vital reagents and analytical tools. JG, KM, JG, JC, PP, AC, TC, CR, MC, SP, MS, NHO, MR, MA and RLL contribute with the recruiting and collection of clinical samples and clinical associated data. TPV, CC, CYG, HW and SW wrote the manuscript. All authors critically read the manuscript.

## Disclosure of Conflicts of Interest

Nothing to declare.

## Abbreviations

CTCs: Circulating tumour cells
Ctrl: Control
EMT: Epithelial–mesenchymal transition
HD: Healthy donor
HD RBCs: RBCs isolated from healthy donors
M0: Non-metastatic
M0 RBCs: RBCs isolated from non-metastatic patients
M1: Metastatic
M1 RBCs: RBCs isolated from metastatic patients
NSCLC: Non-small cell lung cancer
OS: Overall survival
PAKi: p21 activated kinase 4 inhibitor
PFS: Progression-free survival
RBCs: Red blood cells
RDW: Red cell distribution width

## Notes

### Competing Interest Statement

The authors have declared no competing interest.

