## Supplemental data_Red blood cell-tumour cell interactions promote tumour cell progression for "Red blood cell-tumour cell interactions promote tumour cell progression"

### Supplemental Methods:

#### RBC isolation and tumour cell priming

Blood samples were collected in EDTA tubes and RBCs were isolated according to previously established protocols within two hours of collection[1]. RBCs were either used directly or preserved with a glycerol buffer containing glycerol 6.2 M, Lactate 0.14 M, KCl 5 mM and $Na_2PO_4$ 5 mM (Sigma Aldrich, USA) and stored at -80°C until further use[2].

The frozen RBCs were thawed at 37°C in a water bath. Afterwards, RBCs were transferred into a 15 mL tube containing 5 mL of a sterile 12% NaCl solution and centrifuged for 5 min at 200 g. The supernatant with lysed RBCs and glycerol was discarded; the pellet was resuspended in a sterile 3.4% NaCl solution and centrifuged again for 5 min at 200 g. Next, 20 µL of RBC pellet was collected and resuspended in fresh media without FCS.

#### Migration assay

Human cancer cell lines were primed for 24h with RBCs isolated from HD, M0 and M1 blood samples. Non-primed tumour cells were included as a control (n= 5/group, in triplicates). The migration of the primed cells towards an FCS gradient was assessed. In parallel, migration towards a RBCs suspension was also assessed to study an indirect interaction effect of RBCs and tumour cells.

The transwell migration assays using the primed tumour cell lines were performed by adding 200 µL FCS-deprived media containing 50,000 tumour cells into cell culture inserts (BD Falcon, DE; 8 µm pores), while media containing 10% FCS as a chemoattractant was placed in the lower chambers. After 8 h, the media was removed, and the membrane was washed in PBS and fixed with 4% (w/v) PFA/PBS for 10 min. Cells were permeabilized with methanol for 20 min and stained with 0.5% (w/v) crystal violet/$H_2O$ solution for 10 min. The excess staining solution was removed by washing with PBS. Non-migrated cells were scraped off from the inside of the inserts with sterile cotton. The membrane was imaged under a brightfield microscope (Leica DMi8, Leica Microsystems/Axio Observer 7, Zeiss) and cells were counted manually. To study an indirect interaction effect of RBCs on the migration of tumour cells, the transwell migration assay was performed by adding 200 µL FCS-deprived media containing 10,000 cells into cell culture inserts, while media containing 10% FCS and 440,000 RBCs was placed in the lower chambers. After 16 h, the media was removed, and the membrane was washed, fixed and stained as previously described.

**Proliferation assay**

Proliferation of cell lines was assessed by MTT (3-(4,5-dimethylthiazol-2-yl)-2,5-diphenyltetrazolium bromide) assays by indirect measurements of metabolic conversion of MTT to formazan. First, 4,000 tumour cells/well were seeded into 96-well plates in 100 µL medium and 400,000 RBCs were added to each well. Proliferation was measured at 48 and 72 h. For the measurement, 20 µL of MTT was added to the individual wells. After 3 h, the media was carefully removed, and formazan was dissolved in 100 µL of DMSO. Absorption was measured at 540 nm and 650 nm (reference wavelength) using a NanoQuant infinite M200Pro ELISA plate-reader (Tecan, Switzerland) (n= 5/group, sextuplicate).

**Static adhesion to endothelial cells**

The adherence of tumour cells to HUVEC cells was studied by using human cancer cell lines primed with HD or M1 RBCs as well as non-primed control cells. For breast cancer, MDA-MB-231 cells expressing GFP were used, while for NSCLC, H1975 cells were stained with CellTracker™ Green CMFDA Dye (ThermoFisher, USA) following manufacturer's instructions. Briefly, 80,000 HUVEC cells were plated on 12 well plates (VWR, USA) and cultured for 2 days until a monolayer was formed. Tumour cells were seeded on top of the HUVEC monolayer and allowed to attach for 1h at 37°C. After incubation, the media was removed and wells were washed 3 times with PBS. Next, new media was added and cells were imaged using a fluorescence microscope (Leica DMi8, Leica Microsystems/Axio Observer 7, Zeiss). Pictures of 8 random fields of view were taken for each condition and cells were counted with the bioimage analysis software QuPath (v 0.5.1) (n= 5/group, triplicates).

**Adhesion to collagen I**

MDA-MB-231 cells primed with HD, M0 or M1 RBCs, as well as a non-primed control, were seeded on a 48-well Cytoselect Collagen I Plate (Cell Biolabs, USA) and incubated for 1 h at 37 °C and 5 % $CO_2$ – 100,000 cells/condition. After incubation, media was aspirated and wells were washed with PBS. Adhered cells were stained following the manufacturer´s instructions. Absorbance was measured at 560 nm using a BIOTEK EPOCH 2 microplate reader (Agilent Technologies, USA) (n= 5/group, triplicates).

**Electric cell-impedance sensing (ECIS)**

Endothelial cell monolayer impedance was measured continuously using an ECIS 1600R instrument (Applied BioPhysics, Inc.) as previously described[3]. HUVECs were grown to confluence in standard EBM-2 medium supplemented with 10% heat-inactivated foetal calf serum (FCS), 1% non-essential amino acids, 1% glutamic acid, and 1% penicillin/streptomycin on planar gold film electrodes placed at the bottom of an 8-well electrode array (ECIS Cultureware 8W10E+, Applied BioPhysics Inc, Troy, NY, USA) coated with gelatine (0.5% w/v) for 30 min. Measurements were made under standard cell culture conditions. Impedance was measured continuously at different frequencies (500-64,000 Hz). After 24 h, half of the endothelial cell culture medium was replaced by medium containing 10,000 H1975 cells (non-primed and HD, M1 primed RBC) or supernatants (H1975, RBCs from HD and M1). Results are expressed as relative electrical resistance with a 95% confidence interval (95% CI) (n= 4/group, triplicates). Thrombin (1 unit/ml) was used as positive control (Sigma-Aldrich, USA).

#### *In vivo* Zebrafish experiments

Zebrafish embryos (*Danio rerio*, wild type) were generated by natural spawning as previously described[4]. Zebrafish embryos were collected at 0 hpf (hours post fertilization) and incubated at 28.5°C in petri dishes until 48 hpf. Human cancer cell lines were primed 24 h before injection with RBCs isolated from HD or M1 patients. Primed cells were trypsinized and concentrated in an Eppendorf tube at a rate of ≈ 1 million cells in 10 µL of PBS with 2% of PVP (Polyvinylpyrrolidone, Sigma-Aldrich, USA) to avoid cellular aggregation. Cells were stained using Vibrant Dil following the manufacturer’s protocol (ThermoFisher, USA). After cell preparation, 48 hpf embryos were anesthetized with 0.02% of tricaine (Sigma-Aldrich, USA) and cell injection was performed using borosilicate needles (1 mm O.D. x 0.75 mm I.D.; World Precision Instruments). Between 100 and 200 tumour cells were injected into the circulation (Duct of Cuvier) of each fish embryo using a microinjector (IM-31 Electric Microinjector, Narishige, UK) with an output pressure of 20 kPA and 15 ms of injection time per injection. Afterwards, embryos were incubated until 1 day post-injection at 34°C (n= 3/group, 33 fish injected/sample). Imaging of the injected embryos 1 day post-injection was performed using a fluorescence stereomicroscope (AZ-100, Nikon, Japan) to measure the spreading and proliferation of the tumour cells in the caudal hematopoietic tissue in each of the conditions assayed.

#### Immunofluorescence

Samples were fixed with 2% PFA (Sigma-Aldrich, USA) for 10 min. The samples were washed with 0.5 mL of 1x-PBS before permeabilization with 0.1% Triton X 100/PBS (Sigma-Aldrich, USA) for 15 min. Following two additional wash steps, 10% AB-serum/PBS (BioRad, Germany) was applied for blocking (60 min). Actin filaments were stained with Alexa Fluor™ 488 Phalloidin (ThermoFisher, USA) according to the manufacturer's instructions, followed by 5 min of DAPI-incubation (1 µg/mL). K5, CDH1 and VASP were stained with the primary antibodies Mouse Anti-Cytokeratin 5 (clone: XM26) (Abcam, UK), Mouse Anti-E-Cadherin (clone: 36) (BD Bioscience, USA), and Mouse Anti-VASP (clone: A-11) (Santa Cruz, USA). Images were acquired using a fluorescence microscope (Axio Observer 7, Zeiss). Protrusions on the tumour cells were quantified with ImageJ (n= 5/group, triplicates).

**Western blot**

To prepare cell lysates, cell suspensions were collected after splitting and centrifuged at 500×g for 5 min, resuspended with PBS for washing, and centrifuged again. The cell pellet was then dissolved in 200 µL M-PER lysis buffer (ThermoFisher, USA) and frozen at −20°C overnight to disrupt the cells. To remove cell debris, the lysates were thawed on ice, centrifuged at 15,000 ×g at 4°C for 20 min, and the supernatant was collected as the finished cell lysate. Cell lysates were stored at −20°C. For Western blotting, the protein concentration of the lysates was determined using the Bradford assay with Protein Assay Dye Reagent (Bio-Rad, USA) and a BSA standard (ThermoFisher, USA). Samples were prepared with a final protein concentration of 1 µg/µL in SDS sample buffer (Laemmli buffer). Samples were denaturalized at 95°C for 5 min and afterwards stored at −20°C.

Western blotting was performed by standard procedures, using nitrocellulose membranes (ThermoFisher, USA). Antibodies and conditions used are listed in Supplemental Methods. For quantification, blots were performed threefold using three different lysates, imaged using chemiluminescence imagers (Intas ECL Chemocam, Vilber Fusion FX), and expression was analysed using ImageJ (n= 4/group, triplicates).

**Mass spectrometry**

For protein digestion, samples were dissolved in 100 mM triethyl ammonium bicarbonate and 1% w/v sodium deoxycholate buffer, boiled at 95°C for 5 min and sonicated with a probe sonicator. The protein concentration of denatured proteins was determined by the Pierce bicinchoninic acid assay Protein assay kit (ThermoFisher, USA) and samples were diluted to 20 µg of protein in 50 µL buffer. Disulfide bonds were reduced in 10 mM dithiothreitol for 30 min at 56°C and alkylated in the presence of 20 mM iodoacetamide for 30 min at 37°C in the dark. Digestion with trypsin was performed (Porcine, sequencing grade, Promega, USA) at 1:100 (enzyme: protein) ratio at 37°C overnight and digestion was stopped by adding 1% formic acid. Samples were centrifuged at 14 000 g for 5 min and the supernatant was dried in a vacuum centrifuge and stored at -20°C until further use (n= 5/group, 4 replicates).

For the LC-MS/MS acquisition, the dried peptides were dissolved in 0.1% FA to a concentration of 1 µg/µL and 1 µg of peptides was injected into the LC-MS/MS system. Chromatographic separation of peptides was achieved with a two-buffer system (buffer A: 0.1% FA in $H_2O$, buffer B: 0.1% FA in ACN) on a nano-UHPLC (Dionex Ultimate 3000 UHPLC system, ThermoFisher, USA). Attached to the UHPLC was a peptide trap (100 µm x 20 mm, 100 Å pore size, 5 µm particle size, C18, Nano Viper, ThermoFisher, USA) for online desalting and purification, followed by a 25 cm C18 reversed-phase column (75 µm x 250 mm, 130 Å pore size, 1.7 µm particle size, peptide BEH C18, nanoEase, Waters, USA). Peptides were separated using an 80 min method with linearly increasing ACN concentration from 2% to 30% ACN over 60 min.

MS/MS measurements were performed on a quadrupole-orbitrap hybrid mass spectrometer (QExactive, ThermoFisher, USA). Eluting peptides were ionized using a nano-electrospray ionization source (nano-ESI) with a spray voltage of 1,800 and analysed in data-dependent acquisition (DDA) mode. For each MS1 scan, ions were accumulated for a maximum of 240 ms or until a charge density of 1 x $10^6$ ions (AGC Target) was reached. Fourier-transformation-based mass analysis of the data from the orbitrap mass analyser was performed covering a mass range of m/z 400 – 1,200 with a resolution of 70,000 at m/z = 200. Peptides being responsible for the 15 highest signal intensities per precursor scan with a minimum AGC target of 5 x $10^3$ and charge state from +2 to +5 were isolated within a m/z 2 isolation window and fragmented with a normalized collision energy of 25% using higher energy collisional dissociation (HCD). MS2 scanning was performed, covering a mass range starting at m/z 100 and accumulated for 50 ms or to an AGC target of 1 x $10^5$ at a resolution of 17,500 at m/z = 200. Already fragmented peptides were excluded for 20 s.

LC-MS/MS data were searched with the Sequest algorithm integrated into the Proteome Discoverer software (V3.0.0.757, ThermoFisher, USA) against a reviewed human database, obtained in December 2022. The technical replicates were combined into fractions while loading them into the search engine. Carbamidomethylation was set as a fixed modification for cysteine residues. The oxidation of methionine as well as acetylation and loss of methionine at the protein N-terminus were allowed as variable modifications. A maximum number of two missing tryptic cleavages was set. Peptides between 6 and 144 amino acids were considered. A strict cut-off (FDR < 0.01) was set for peptide and protein identification. Quantification was performed using the Minora Algorithm, implemented in Proteome Discoverer. For LS/MS data analysis, protein abundances were loaded into Perseus (Max Plank Institute for Biochemistry, Version 2.0.3[5]) and log2 transformed to approximate the Gaussian probability distribution. Protein abundances were median normalized by column to cope for injection-related discrepancies in total protein amounts. Data was reduced to valid values only, for principal component analysis. Visualization was performed in GraphPad Prism. Student's T-testing was performed based on proteins quantified in ≥ 3 samples per biological phenotype with a P-value cut-off of $P < 0.05$ and a false discovery rate (FDR) of $q < 0.05$. *T*-testing results were visualized using an in house-script, implementing the ggplot2 and ggrepel packages in the R-software environment (version 4.4.2). Based on the FDR-significant and 2-fold changed proteins, IPA (Ingenuity Pathway Analysis, Qiagen)[6] was performed. Canonical pathways analysis identified the pathways from the QIAGEN IPA library of canonical pathways that were most significant to the data set. Molecules from the data set that met the significance cut-off (q-value < 0.05, fold-change > 2) and were associated with a canonical pathway in the QIAGEN Knowledge Base were considered for the analysis. A right-tailed Fisher's Exact Test was used to calculate a p-value determining the probability that the association between the genes in the dataset and the canonical pathway is explained by chance alone (P-value < 0.05). A z-score was calculated to indicate the likelihood of activation or inhibition of that pathway.

### Supplemental Figures:

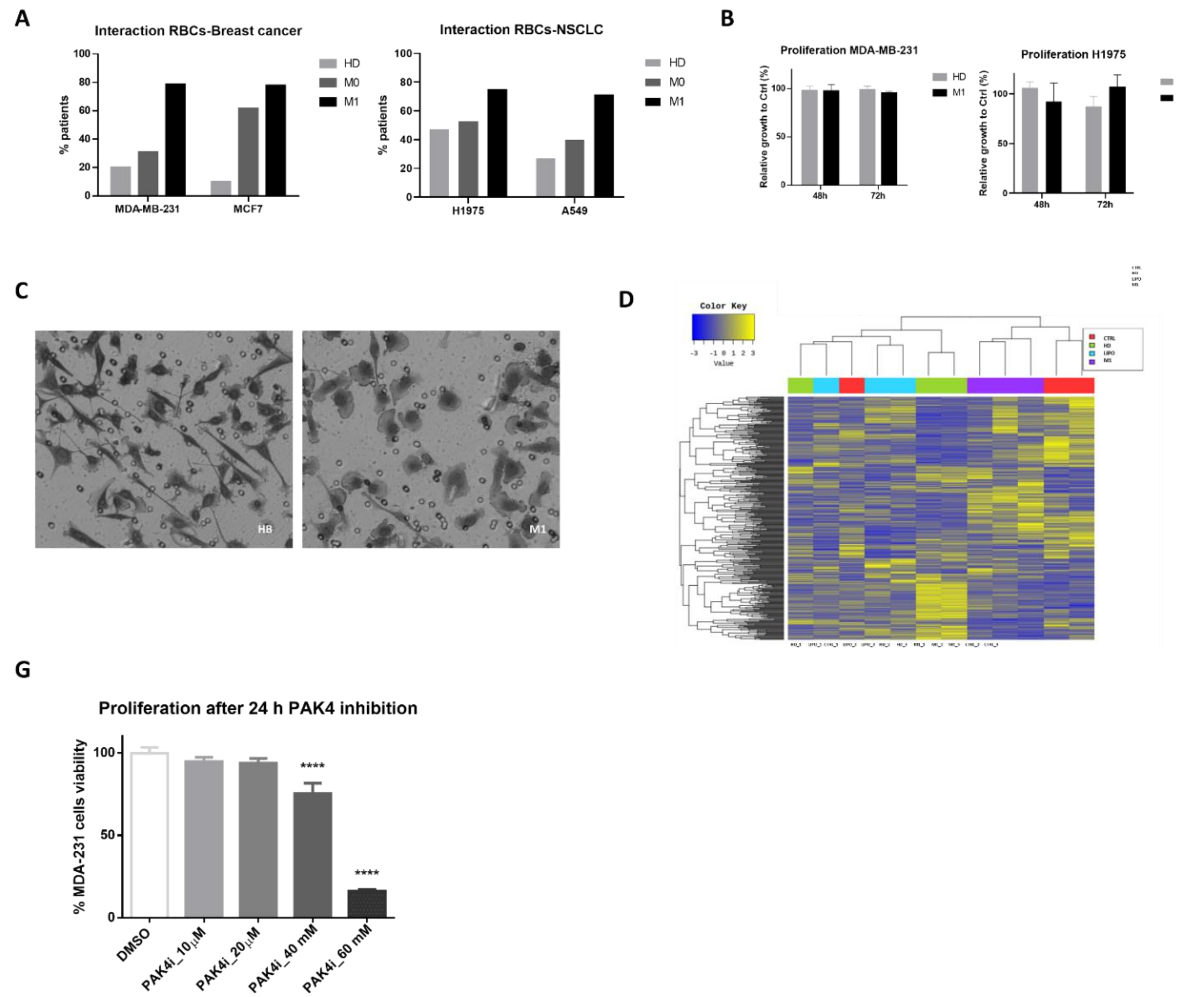


**Supplemental Figure 1. (A)** Graph representing the percentage of cancer patients (M0 and M1) or HD whose RBCs interact *in vitro* with either breast cancer (MDA-MB-231 and MCF7) or NSCLC (H1975 and A549) cells from our clinical cohort (n=15/group, triplicates). **(B)** Proliferation assay (MTT) of breast cancer (MDA-MB-231) or NSCLC (H1975) cells measured after 48 and 72 h of co-culture with M1 or HD RBCs, compared with the non-co-cultured cells rate of growth (Ctrl). **(C)** Representative images of MDA-MB-231 observed in migrated cells from indirect transwell assay after priming with HD (left panel) and M1 (right panel), this latter depicting morphological changes. **(D)** Heatmap showing results of the hierarchical clustering analysis of MDA-MB-231 (Ctrl, red), primed with HD (green), M1 (purple) or Liposomes (LIPO, blue), n=3. **(E)** Graph representing the cell viability of MDA-MB-231 after 24 h exposure to different doses of the PAK4 inhibitor by MTT assay. 20 µM was the selected concentration for inhibitory assays since does not impact cellular proliferation. Since PAK4i is diluted in DMSO, the same volume of DMSO was also tested to rule out any effects due to DMSO toxicity. * P < 0.05, ** P < 0.01, *** P < 0.001.

A

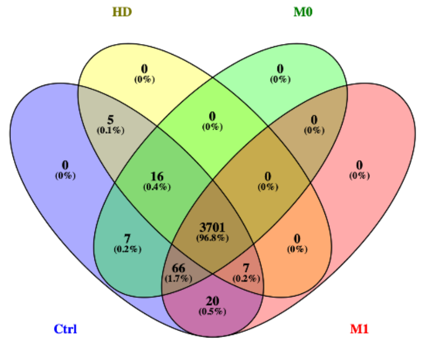


B

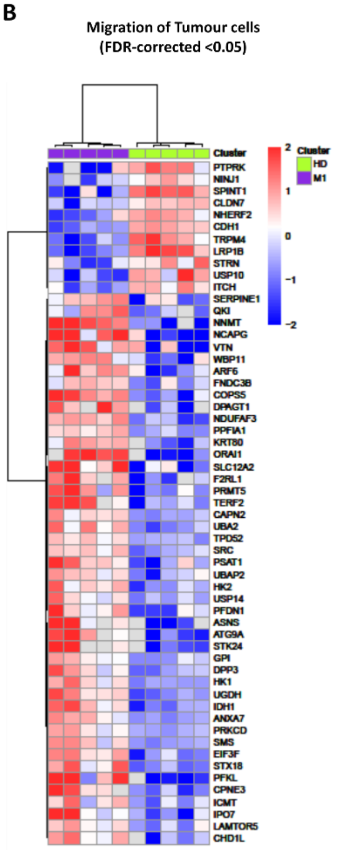


C

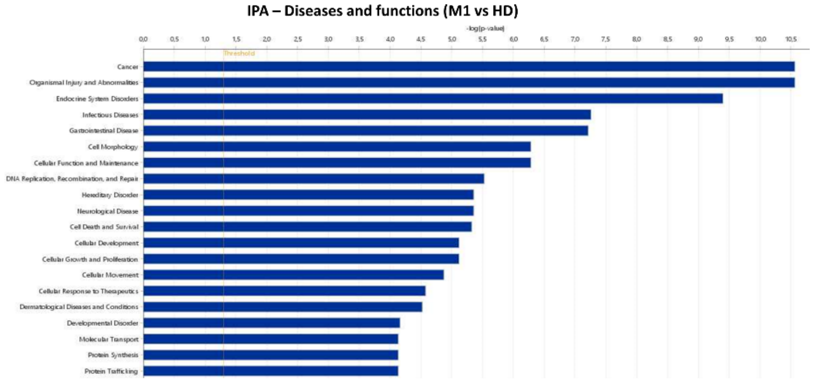


D

| Upstream Regulator | Molecule Type | Predicted Activation State | Activation z-score | p-value of overlap | Target Molecules in Dataset |
|---|---|---|---|---|---|
| PCGEM1 | other | Activated | 2,060 | 1,85E-05 | G6PD,GPI,HK2,IDH1,PGD,PGLS,TALDO1 |
| IL15 | cytokine | Activated | 3,293 | 2,43E-05 | ALDOC,G6PD,GPI,HK1,HK2,PFKL,PGD,PGM1,PSMB9,RBX1,TALDO1 |
| ERBB2 | kinase | Activated | 2,168 | 3,45E-03 | APLP2,CDH1,CHD1L,DAAM1,EPHB2,GINS3,GNPTAB,JAG1,MCM3,NCAPG,PRSS3 ,PTPRK,RFC3,RPA1,RPA2,RRM2,SERPINE1 |
| TFEB | ranscription regulato | Activated | 2,529 | 5,82E-03 | CDH1,FOSL1,JAG1,MCM3,NCAPG,NUP160,NUP188,NUP54,RFC3,RPA2,RRM2 |
| BCR (complex) | complex | Activated | 3,000 | 8,45E-03 | ASNS,ATP6V1G1,GFPT1,GPI,HK1,HK2,PPA1,SRM,TMEM97 |
| RABL6 | other | Activated | 2,236 | 1,05E-02 | DUT,NCAPG,PSME3,RFC3,TMEM97 |
| EGF | growth factor | Activated | 2,156 | 2,47E-02 | CDH1,PSMB1,PSMB3,PSMB4,SERPINE1 |
| HSF1 | ranscription regulato | Activated | 2,219 | 7,27E-02 | ANXA11,ATAD2,CDH1,CLDN7,MCM3,RANBP1 |
| LDHB | enzyme | Inhibited | -2,000 | 3,93E-04 | ALDOC,GPI,HK1,HK2 |
| TGFB2 | growth factor | Inhibited | -2,414 | 3,79E-03 | ASNS,HK2,JAG1,PGM1,PSAT1,SERPINE1,SRM,VARS1 |
| RAD51 | enzyme | Inhibited | -2,000 | 4,79E-02 | PSMC6,PSMD11,PSMD14,PSME3 |

E

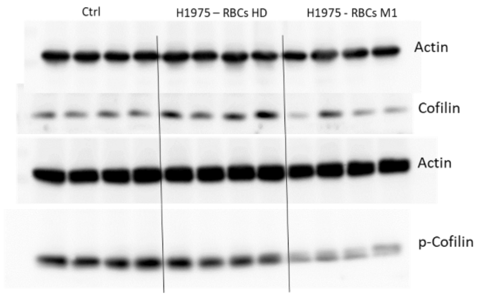


F

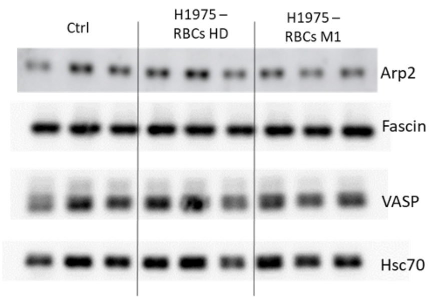

**Supplemental Figure 2. (A)** Venn-diagram representing all the proteins quantified in all the samples included on the LC-MS/MS method by DIA. **(B)** Hierarchical clustering of the differential expression of the proteins from the dataset involved in migration of cancer cell lines, between H1975 cells primed wither with HD or M1 RBCs (FDR < 0.05, Fold-change >2, P = 0.0103). **(C)** Top 20 diseases and functions predicted to be affected with IPA analysis (z-score > 2). **(D)** Upstream regulatory proteins were predicted to be activated and inhibited with IPA analysis (z-score > 2 and > -2, respectively). **(E)** Protein levels of Cofilin and p-Cofilin were analysed by western blot from H1975 cells primed with M1 and HD, as well as non-primed cells (Crtl) (n= 4/group). **(F)** Protein levels of Arp2, VASP and Fascin were analysed by western blot from H1975 cells primed with M1 and HD, as well as non-primed cells (Crtl) (n= 3/group).

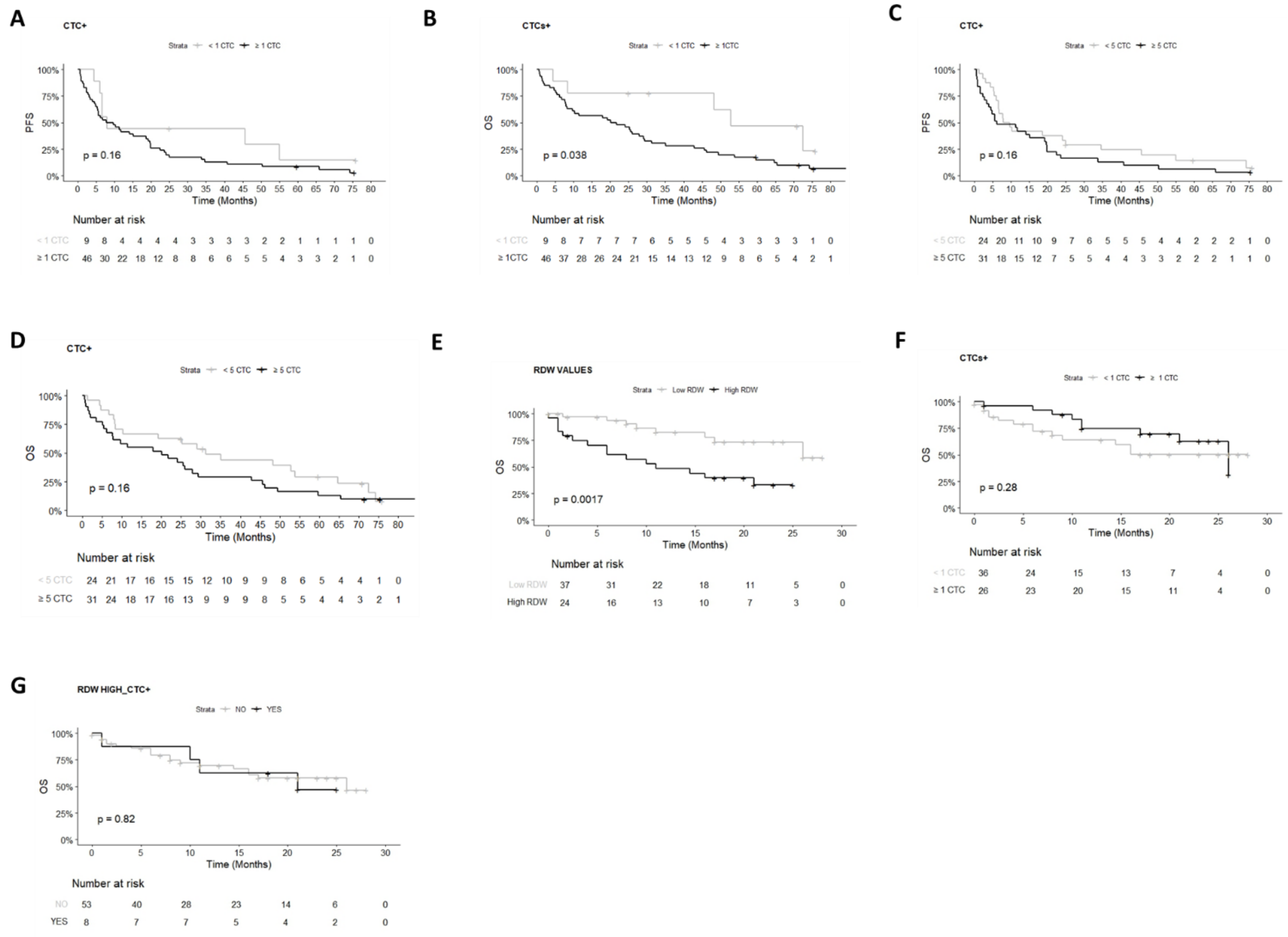


**Supplemental Figure 3. (A-B)** Kaplan-Meier analysis of the presence of ≥ 1 CTC/7.5ml blood (CellSearch®) (black) or < 1 CTC (grey) in metastatic breast cancer patients (n= 55) for PFS: 7.99 vs 8.75 months, P = 0.2 **(A)** or OS: 52.8 vs 20.9 months, P = 0.04 **(B). (C-D)** Kaplan-Meier analysis of the presence of ≥ 5 CTC (black) or < 5 CTC (grey) (n=55) in breast cancer patients for PFS: 8.86 vs 6.21 months, P = 0.2) **(C)** or OS: 31.2 vs 20.1 months, P = 0.2 **(D). (E)** RDW values (high- black- or low- grey-; cut-off = 14.5) in NSCLC (n=61) (undefined vs 11 months, P = 0.0017). **(F)** The presence of ≥ 1 CTC (black) or < 1 CTC (grey) by Parsortix® from 7.5 mL of blood from NSCLC patients (n=61) for OS: undefined vs 26 months, P = 0.28. **(G)** Combination of high RDW and ≥ 1 CTC (RDW HIGH_CTC+, black) or not (grey) (RDW cut-off = 14.5) (n=61) OS: 26 vs 21 months, P = 0.82. P-values were calculated using the log-rank test.

### Supplemental Tables:

**Table 1. Clinical characteristics of the cohort of breast cancer and HD samples** used for RBCs isolation and further RBCs experiments.

| Category | M1 | | M0 | | HD | |
|---|---|---|---|---|---|---|
| | Media | SD | Media | SD | Media | SD |
| Age (years) | 62.76 | 12.22 | 54.07 | 9.863 | 57.55 | 9.12 |
| Tumour stage | n | % | n | % | n | % |
| 0 | | | | | 29 | 100 |
| I-III | | | 29 | 100 | | |
| IV | 33 | 100 | | | | |
| Subtype | | | | | | |
| Luminal | 20 | 60.6 | 21 | 72.5 | | |
| Her2 | 1 | 3 | 5 | 17.2 | | |
| Triple Negative | 6 | 18.2 | 3 | 10.3 | | |
| Mixed subtypes | 6 | 18.2 | | | | |
| Metastasis Location | | | | | | |
| Bone | 7 | 39.4 | | | | |
| Visceral | 12 | 21.2 | | | | |
| Bone & Visceral | 13 | 36.4 | | | | |
| Others | 1 | 3 | | | | |

**Table 2. Clinical characteristics of the cohort of lung cancer (NSCLC) and HD samples** used for RBCs isolation and further RBCs experiments.

| Category | M1 | | M0 | | HD | |
|---|---|---|---|---|---|---|
| | Media | SD | Media | SD | Media | SD |
| Age (years) | 66 | 10.1 | 70.9 | 8.1 | 57.4 | 6.1 |
| Tumour stage | n | % | n | % | n | % |
| 0 | | | | | 25 | 100 |
| I-III | | | 17 | 100 | | |
| IV | 41 | 100 | | | | |
| Histology (NSCLC) | | | | | | |
| ADCA | 34 | 82.9 | 9 | 53 | | |
| SqC | 4 | 9.8 | 8 | 47 | | |
| Unknown | 3 | 7.2 | 0 | 0 | | |

ADAC: adenocarcinoma, SqC= squamous cell carcinoma

**Table 3**. RT-qPCR Taqman probe information.

| Gene | Taqman Reference |
|---|---|
| B2M | Hs00187842_m1 |
| PAK4 | Hs01100061_m1 |
| PLS3 | Hs00543971_m1 |
| VIM | Hs00958116_m1 |

**Table 4. RNAseq analysis of MDA-MB-231 cells**. Top 20 genes with higher fold-change difference in the metastatic group compared with healthy donors.

| Gene | baseMean | log2FoldChange | lfcSE | stat | p-value | p-adj |
|---|---|---|---|---|---|---|
| TRNY | 204,52 | -3,12 | 0,30 | -5,42 | 6,11E-08 | 0,00022 |
| RNR1 | 24,55 | -22,99 | 0,95 | -4,78 | 1,77E-06 | NA |
| ND1 | 94,38 | -5,82 | 0,53 | -4,77 | 1,86E-06 | 0,002378 |
| ND5 | 227,34 | -4,32 | 0,45 | -4,74 | 2,18E-06 | 0,002525 |
| MIR663A | 244,87 | 2,08 | 0,24 | 4,43 | 9,41E-06 | 0,006638 |
| ITPKB | 120,67 | 2,36 | 0,30 | 4,14 | 3,51E-05 | 0,01373 |
| GOLGA8B | 284,93 | -2,06 | 0,26 | -4,04 | 5,46E-05 | 0,015512 |
| TYMP | 208,80 | 2,13 | 0,27 | 4,01 | 6,06E-05 | 0,016495 |
| TRNC | 100,95 | -2,81 | 0,37 | -4,01 | 6,19E-05 | 0,016498 |
| RNR2 | 171,40 | -2,82 | 0,38 | -3,89 | 0,000102 | 0,023243 |
| ND4 | 184,30 | -3,56 | 0,47 | -3,87 | 0,000109 | 0,024469 |
| IER5L | 457,34 | 2,10 | 0,28 | 3,85 | 0,000119 | 0,025005 |
| SFN | 294,86 | 2,04 | 0,27 | 3,85 | 0,000119 | 0,025005 |
| PDE4A | 163,41 | 2,10 | 0,28 | 3,82 | 0,000133 | 0,026929 |
| ADAT3 | 52,78 | 3,65 | 0,49 | 3,81 | 0,000141 | 0,027394 |
| LINC00996 | 89,34 | 5,25 | 0,63 | 3,78 | 0,000157 | 0,028973 |
| EPHX1 | 60,34 | 3,37 | 0,49 | 3,54 | 0,000398 | 0,047959 |
| RHBDD2 | 116,66 | 2,32 | 0,35 | 3,46 | 0,000533 | 0,05778 |
| SNORD22 | 168,59 | -3,05 | 0,46 | -3,46 | 0,000541 | 0,057891 |
| PAK4 | 181,69 | 2,13 | 0,32 | 3,38 | 0,000735 | 0,069134 |

**Table 5. Proteomic analysis of H1975 cells.** Top 20 genes with higher fold-change difference in the metastatic group compared with healthy donors.

| Protein Information | | | Student's T-test results | | |
|---|---|---|---|---|---|
| Accession | Description | Gene | p-value | q-value | Difference |
| P13647 | Keratin, type II cytoskeletal 5 [OS=Homo sapiens] | KRT5 | 0,003064 | 0,012303 | 5,40072 |
| P40938 | Replication factor C subunit 3 [OS=Homo sapiens] | RFC3 | 0,000729 | 0,006764 | 5,3677 |
| Q16774 | Guanylate kinase [OS=Homo sapiens] | GUK1 | 0,003173 | 0,012518 | 4,55753 |
| Q9BU76 | Multiple myeloma tumor-associated protein 2 [OS=Homo sapiens] | MMTAG2 | 0,007163 | 0,020581 | 4,52277 |
| O43657 | Tetraspanin-6 [OS=Homo sapiens] | TSPAN6 | 0,000464 | 0,005705 | 4,41667 |
| P04732 | Metallothionein-1E [OS=Homo sapiens] | MT1E | 0,024619 | 0,043191 | 4,39436 |
| Q9BRX5 | DNA replication complex GINS protein PSF3 [OS=Homo sapiens] | GINS3 | 0,008092 | 0,022494 | 4,36068 |
| Q9BRF8 | Serine/threonine-protein phosphatase CPPED1 [OS=Homo sapiens] | CPPED1 | 0,004081 | 0,014619 | 4,34866 |
| P23743 | Diacylglycerol kinase alpha [OS=Homo sapiens] | DGKA | 0,011058 | 0,02656 | 4,33836 |
| P60002 | Transcription elongation factor 1 homolog [OS=Homo sapiens] | ELOF1 | 4,04E-05 | 0,001286 | 4,2764 |
| Q5QNW6 | Histone H2B type 2-F [OS=Homo sapiens] | H2BC18 | 9,21E-07 | 0 | -4,42887 |
| Q8NAV1 | Pre-mRNA-splicing factor 38A [OS=Homo sapiens] | PRPF38A | 8,66E-06 | 0 | -1,0376 |
| Q9Y2Z4 | Tyrosine--tRNA ligase, mitochondrial [OS=Homo sapiens] | YARS2 | 1,34E-05 | 0 | -1,03604 |
| Q13641 | Trophoblast glycoprotein [OS=Homo sapiens] | TPBG | 2,69E-05 | 0,000909 | -2,94235 |
| Q8NEY8 | Periphilin-1 [OS=Homo sapiens] | PPHLN1 | 2,67E-05 | 0,000952 | -1,00891 |
| Q92804 | TATA-binding protein-associated factor 2N [OS=Homo sapiens] | TAF15 | 1,72E-05 | 0,001067 | -1,19153 |
| P12830 | Cadherin-1 [OS=Homo sapiens] | CDH1 | 2,33E-05 | 0,001111 | -1,78083 |
| Q92934 | Bcl2-associated agonist of cell death [OS=Homo sapiens] | BAD | 1,54E-05 | 0,001455 | -1,43605 |
| Q9NWT1 | p21-activated protein kinase-interacting protein 1 [OS=Homo sapiens] | PAK1IP1 | 5,45E-05 | 0,001829 | -1,21247 |
| Q16629 | Serine/arginine-rich splicing factor 7 [OS=Homo sapiens] | SRSF7 | 0,000149 | 0,003016 | -1,02059 |
